# Integrative analysis of GTEx data reveals that systemic transcriptomes correlate with human spermatogenic dysfunction

**DOI:** 10.1101/2025.11.19.689370

**Authors:** Xing An, Feng-Yun Xie, Juan Chen, Yingyu Chen, Jun-Yu Ma

## Abstract

Declining of male fertility may stem from intrinsic testicular dysfunction or systemic influences from other tissues. However, beyond the hypothalamic-pituitary-testis axis, little is known about which organs interact with the testes or whether they affect spermatogenic capacity. To investigate this, we analyzed GTEx open-access data, comparing transcriptomic profiles across various tissues in men with high and low testicular spermatogenic activity. Our findings reveal significant associations between testicular function and the pituitary gland, adrenal gland, lungs, liver, and key brain regions involved in neuroendocrine regulation, including the amygdala, substantia nigra, anterior cingulate cortex, and frontal cortex. These results suggest potential extragonadal pathways that modulate male fertility, highlighting candidate biomarkers for testicular dysfunction and novel therapeutic targets for infertility intervention.

## Introduction

Human spermatogenic capacity exhibits significant interindividual variability, and impaired spermatogenesis, resulting from genetic and environmental factors, can lead to reduced male fertility [1–3]. It is well established that testicular function can be disrupted by various pathological conditions, including obesity, diabetes, and inflammation, which exert their effects through metabolic, immune, and endocrine dysregulation [4–6]. However, little is known about the specific organs or tissues in the human body that are most closely associated with testicular spermatogenesis.

From an endocrine perspective, testicular function is regulated by the hypothalamic-pituitary-testicular (HPT) axis [7]. The hypothalamus secretes gonadotropin-releasing hormone (GnRH), which stimulates the pituitary gland to release luteinizing hormone (LH) and follicle-stimulating hormone (FSH). LH promotes testosterone production in Leydig cells, while FSH activates Sertoli cells to facilitate sperm maturation. Testosterone exerts negative feedback on the secretion of GnRH and LH, whereas inhibin, produced by Sertoli cells, suppresses FSH levels. Together, testosterone and FSH works synergistically to regulate spermatogenesis. Despite this complex feedback network, it remains unclear whether the efficiency of testicular spermatogenesis correlates with global gene expression patterns in the hypothalamus and pituitary.

Psychological stress has been identified as a significant modulator of male fertility [8], yet the transcriptional mechanisms that connect mental health to spermatogenic dysfunction remain poorly understood. Emerging evidence suggests a bidirectional relationship between brain function and testicular activity. For example, testosterone levels influence the development and reactivity of brain regions such as the hippocampus and amygdala [9–11]. Notably, the hippocampus can regulate testosterone synthesis independently of the pituitary pathway [12]. Furthermore, comparative transcriptomic analyses reveal unexpected similarities in gene expression between the brain and testes [13]. Collectively, these findings highlight the importance of investigating brain-testicular interactions, as they may have implications for both neurological disorders and male reproductive health.

In humans, the adrenal gland plays a crucial role in regulating stress responses, metabolism, and blood pressure, while also producing minor sex hormones such as dehydroepiandrosterone. Similarly, the thyroid gland modulates metabolism, growth, and development through the secretion of thyroid hormones (T3 and T4), which exert systemic effects, including influences on reproductive function. Although both the adrenal and thyroid glands are known to interact with testicular function, it remains unclear whether their transcriptional dysregulation is linked to variations in spermatogenic capacity.

The Genotype-Tissue Expression database (GTEx) provides a comprehensive repository of transcriptomic data derived from various tissues, including the testis, across a large human cohort [14]. Our previous research has established that testicular transcriptome data from GTEx reliably capture interindividual differences in spermatogenic efficiency. Utilizing this resource, we conducted a natural experiment to examine transcriptomic variations across multiple tissues in individuals stratified by their spermatogenic capacity. Specifically, we compared gene expression profiles in brain tissues, HPT axis, adrenal-thyroid systems, and other organ systems between groups with high and low spermatogenic potential. This study elucidates the transcriptional correlates of spermatogenic capacity in non-testicular tissues, providing novel insights into the systemic regulation of male fertility. Our findings may pave the way for improved diagnostic strategies for male infertility and may identify new novel therapeutic targets for male subfertility.

## Materials and methods

### Ethics approval

This study was approved by the Ethic committee of Guangdong Second Provincial General Hospital (Approval No: 2025-KY-KZ-215-01). Clinical trial number: not applicable.

### Data collection from GTEx

In this study, all analyses were conducted using publicly accessible data from GTEx. This dataset includes RNA sequencing data from various tissues, as well as metadata for each donor subject, which were downloaded from https://www.gtexportal.org/ (Table S1). The GTEx Analysis V10 version was utilized for this research. The gene expression data and metadata were converted into AnnData format (version: 0.11.3)[15] using Jupyter Notebook (https://jupyter.org/).

### Testis clustering

Testis clustering was conducted using the single cell RNA-seq analysis tool Scanpy (version 1.10.4) [16]. The gene expression values for each testis, reported as TPM (transcripts per million), were normalized using the sc.pp.normalize_total() function and transformed to log(TPM + 1) using the sc.pp.log1p() method. The ranked genes for each testis cluster were identified using the Wilcoxon method. Additional methods for testis clustering were executed with the default parameters in Scanpy.

### Differential gene expression screening and gene set enrichment analysis

The differentially expressed genes were screened using Diffxpy (v0.7.4, https://github.com/theislab/diffxpy). Gene set enrichment analysis in this study was conducted using ShinyGO (version 0.82) [17]. The genes exhibiting the most significant differential expression, particularly those involved in critical biological pathways such as cellular metabolism, neuroendocrine regulation, and inflammatory responses, were selected for analysis and presentation.

### Statistics analysis

The de.test.t_test() method in Diffxpy, which performs Welch’s t-test for genes expressed in two groups, was utilized for screening differentially expressed genes in testicular data. When the testicular data were analyzed alongside other tissues, as the donor subject number reduced, the de.test.two_sample() method in Diffxpy with the parameter test=’rank’ was used, which use Wilcoxon rank sum (Mann-Whitney U) test to identify the differentially expressed genes between the two groups.

## Results

### GTEx data collection and testis clustering

In this study, testicular RNA-seq data from GTEx were collected, transformed into an AnnData format and analyzed using Scanpy. As a result, 414 testes were clustered into six distinct groups, labeled T1 through T6. Within these testicular clusters, T5 and T6 exhibited high expression levels of genes such as the transcription regulator *CRTC1* and the metabolic regulator *ACOT9*. In contrast, clusters T1, T2, T3, and T4 showed elevated expression of genes like *PCYT2* and *S1PR2*. Notably, the *SLC9C2* gene was highly expressed in T3, while the *TDRD15* gene was highly expressed in T4 (**Figure 1A**). Then we analyzed the gene marker of somatic cells (*VIM* marked all somatic cells, *CD14* marked macrophage cells, *VWF* marked endothelial cells, *DLK1* marked Leydig cells, and *ACTA2* marked myoid cells) and maturing sperm (maker genes include *HOOK1*, *TNP2*, *PRM3*, *SPATA18*, *SPATA7* and *SPEM3*) [18] in these testis clusters, and found that somatic cell marker genes were highly expressed in T5 and T6, whereas mature sperm marker genes were highly expressed in T1 and T2 (**Figure 1B**). High expression of mature sperm genes in T1 and T2 indicates that the cell ratio of sperm cells in testes is high. In contrast, in T5 and T6 testis clusters, high expression of somatic cell markers indicates a high ratio of somatic cells in these testes.

**Figure 1.**
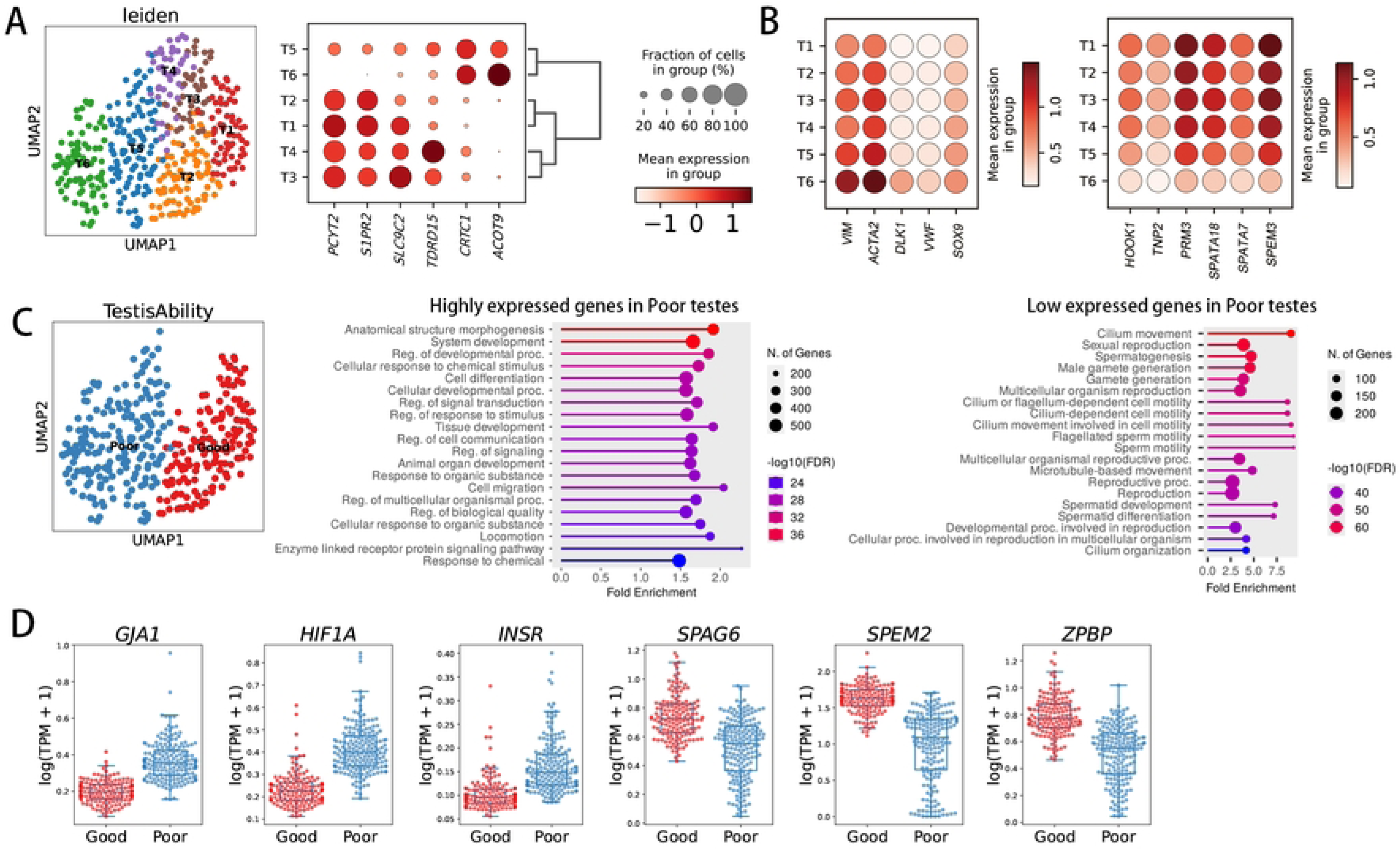
Classification of spermatogenic capacity based on testicular transcriptomes. (**A**) Clustering of testes according to their gene expression profiles. (**B**) The expression patterns of somatic cell marker genes and sperm marker genes in the cell clusters. (**C**) Gene set enrichment analysis of differentially expressed genes in testes exhibiting reduced spermatogenic ability. The testes were categorized into ’good’ and ’poor’ groups based on their spermatogenic capacity. (**D**) Genes that are either upregulated or downregulated in testes with diminished spermatogenic ability.

To facilitate the analysis of testicular transcriptome data, we combined clusters T1 and T2 into the ’Good’ group, representing testes with superior spermatogenic capability, and merged clusters T5 and T6 into the ’Poor’ group, representing testes with diminished spermatogenic capability (**Figure 1C**). We then screened for differentially expressed genes between the Good and Poor testes using Diffxpy. As a result, 1846 genes were found to be highly expressed in the Poor group testes, while 2190 genes exhibited low expression in this group (**Dataset S1**). The highly expressed genes in Poor testes were primarily enriched in pathways related to cellular responses to chemical stimulus and regulation of cell communication, whereas the low expressed genes in Poor testes were enriched in pathways associated with male gamete generation and sperm motility (**Figure 1C**). Notably, among these differentially expressed genes, we identified genes associated with cell communication and chemical response, such as *GJA1*, *HIF1A*, and *INSR*, which were highly expressed in Poor testes. Conversely, genes associated with sperm maturation, such as *SPAG6*, *SPEM2* and *ZPBP* were expressed at lower levels in Poor testes (**Figure 1D**). These differentially expressed genes may not be attributed to the upregulation or downregulation in specific cell types but rather to changes in the cell counts of particular cell populations.

### Hypothalamus-pituitary-testis transcriptional relationships

To analyze the changes in gene expression in the hypothalamus and pituitary gland between the Good Testis (T.good) group and the Poor Testis (T.poor) group, we extracted RNA-seq data from human subjects whose testis, hypothalamus, and pituitary were all sequenced. A total of 83 subjects were selected, comprising 45 individuals in the T.good group and 38 in the T.poor group (**Dataset S2**). Using the DE gene screening criteria of p-value < 0.01 and mean log2(TPM + 1) > 0.01, we identified 20567 DE genes in the testes, 324 DE genes in the pituitary and 74 DE genes in the hypothalamus (**Figure 2A** and **Dataset S3**). The DE genes in the pituitary were primarily enriched in pathways related to stress response to copper ions and canonical glycolysis (**Figure 2B**). In contrast, the DE genes in the hypothalamus were mainly enriched in pathways such as cellular response to cadmium ions and thyroid hormone transport (**Figure 2C**). Notably, the pituitary exhibited approximately 4.4 times more DE genes than the hypothalamus, indicating a stronger transcriptional association with testicular function.

**Figure 2.**
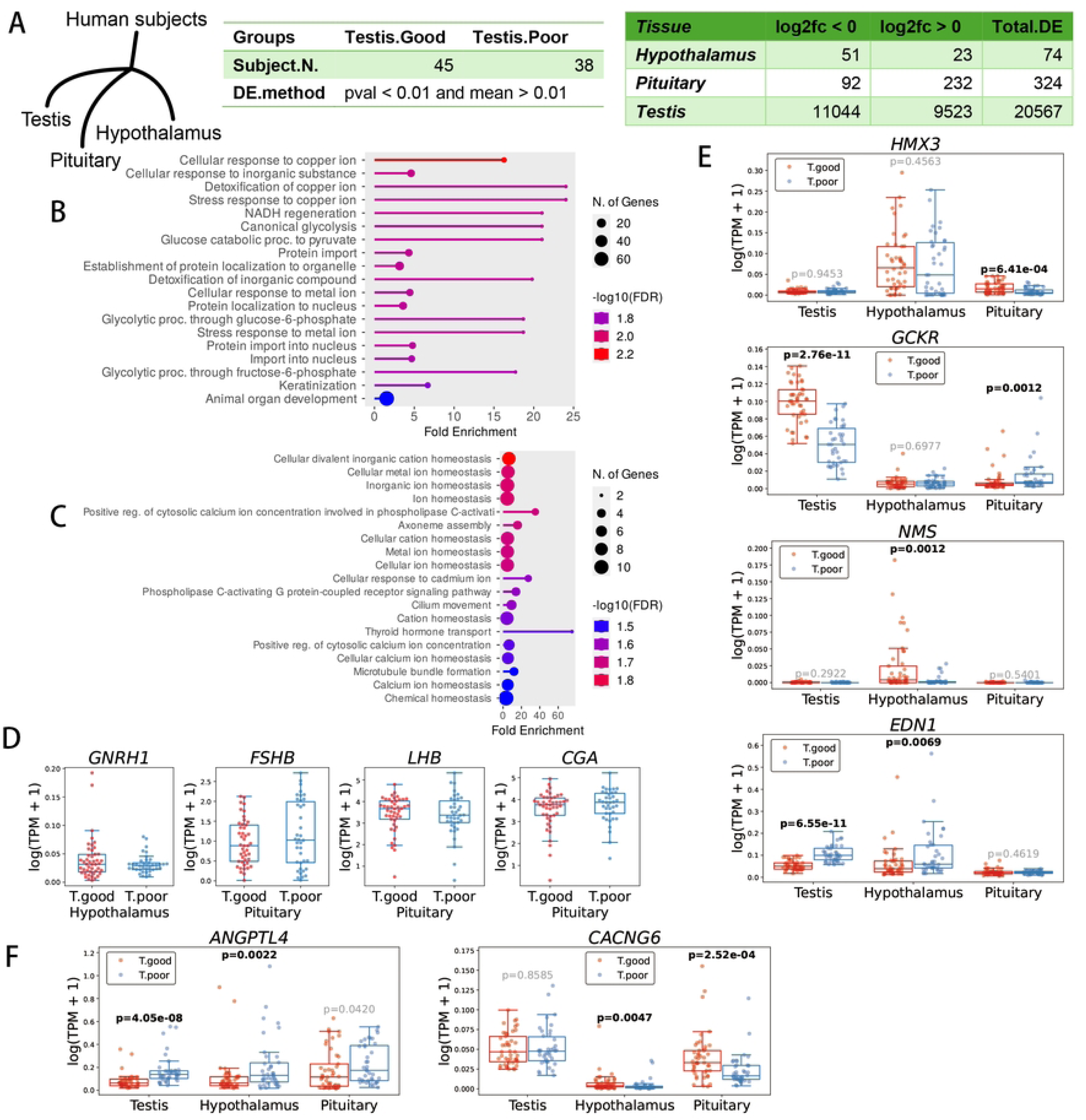
Testis-associated genes in the pituitary and hypothalamus. (**A**) Differentially expressed (DE) genes in the pituitary and hypothalamus of subjects with varying testicular spermatogenesis abilities. (**B**) Pathways enriched by DE genes in the pituitary. (**C**) Pathways enriched by DE genes in the hypothalamus. (**D**) No significant changes in the coding genes for GnRH, LH and FSH in the hypothalamus and pituitary of subjects with poorer testicular function. (**E**) Genes that are differentially expressed in either the pituitary or hypothalamus. (**F**) Genes that are differentially expressed in both the pituitary and hypothalamus.

Within the HPT axis, hypothalamic GnRH and pituitary FSH/LH play critical endocrine roles in regulating testicular spermatogenesis. Interestingly, our analysis revealed no significant expression change in the core neuroendocrine genes *GNRH1* (hypothalamus) or *FSHB*/*LHB*/*CGA* (pituitary) between the T.good and T.poor groups (**Figure 2D**), suggesting that spermatogenic impairment may involve alternative regulatory mechanisms. The homeobox protein gene *HMX3* is essential for the development of the human inner ear and the hypothalamic-pituitary axis [19]. The expression of glucokinase in pituitary may function as a glucose sensor to regulate pituitary secretion [20], while the glucokinase regulatory protein encoded by *GCKR* can inhibit the activity of glucokinase [21]. In the T.poor group, the *HMX3* gene was found to be downregulated in the pituitary, whereas the *GCKR* gene was upregulated (**Figure 2E**). Neuromedin-S peptides are believed to be involved in the regulation of the hypothalamic-pituitary-adrenal axis [22]. The vasoconstrictor endothelin-1, expressed in the hypothalamus, can stimulate the secretion of GnRH [23]. In subjects from the T.poor group, the neuromedin-S coding gene *NMS* was downregulated, while the endothelin-1 coding gene *EDN1* was upregulated in the hypothalamus (**Figure 2E**). In the brain, the angiopoietin-like 4 protein, encoded by *ANGPTL4*, primarily functions in metabolic crosstalk between glia cells and neurons [24]. The voltage-dependent calcium channel gamma coding genes *CACNG1*-*8* have been found associated with schizophrenia [25]. In the T.poor group, the *ANGPTL4* gene was found to be upregulated in both the hypothalamus (p < 0.01) and pituitary (p < 0.05), whereas *CACNG6* was downregulated (**Figure 2F**). All these data indicate that the metabolism and secretion of the hypothalamus and pituitary in T.poor human subjects have been altered.

### Transcriptional associations between the adrenal and thyroid glands and testicular function

Beyond the HPT axis, we investigated two other major endocrine regulators, the adrenal and thyroid glands, for their potential associations with testicular function. From the GTEx database, we identified 85 subjects with available RNA-seq data for all three tissues (testes, adrenal, and thyroid glands), which included 36 individuals with high spermatogenic capacity (T.good) and 59 with impaired spermatogenesis (T.poor) (**Dataset S2**). Using stringent differential expression criteria (q-value < 0.05; mean log2(TPM + 1) >0.01), we observed distinct tissue-specific patterns: testes exhibited extensive transcriptomic alterations (20,972 DE genes), adrenal glands showed moderate changes (260 DE genes) and thyroid glands demonstrated minimal variation (21 DE genes) (**Figure 3A** and **Dataset S4**). The magnitude of transcriptional changes revealed a striking hierarchy of endocrine-testis associations, with adrenal glands exhibiting 12-fold more DE genes than thyroid glands. This quantitative disparity suggests that adrenal function may be more sensitive to, or influential upon, testicular status compared to thyroid activity.

**Figure 3.**
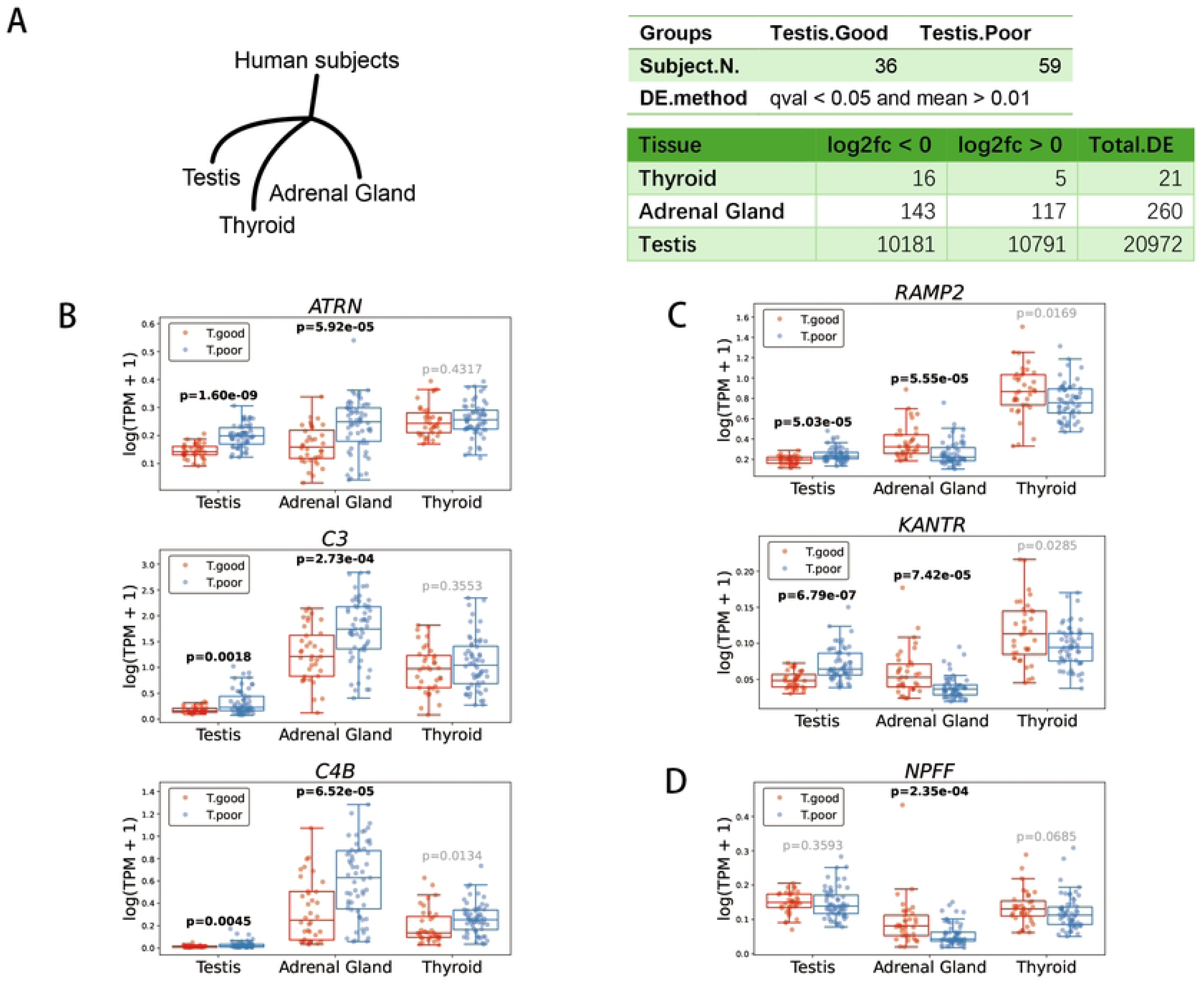
Differentially expressed genes in the adrenal and thyroid glands associated with testicular function. (**A**) Differentially expressed (DE) genes in the adrenal and thyroid glands of subjects with varing testicular spermatogenic abilities. (**B**) Immune response genes were upregulated in the testis, adrenal gland, and thyroid of subjects in the T.poor group. (**C**) Genes that were downregulated in both the adrenal gland and thyroid but upregulated in the testis. (**D**) The gene *NPFF* was downregulated in both the adrenal gland and thyroid, with no significant change observed in the testis.

In the T.poor group, we found the immune response-related genes such as the attractin coding gene *ATRN* and the C3/C4 complement encoding genes *C3* and *C4B*, were upregulated in the adrenal gland. Additionally, only *C4B* was upregulated in the thyroid gland (**Figure 3B**). This suggests that increased inflammatory activity in both adrenal and thyroid tissues may be linked to impaired spermatogenic function in the testes. Among the DE genes, we identified several genes exhibiting consistent expression patterns between the adrenal and thyroid glands. For instance, the *RAMP2* gene, which encodes a protein that regulates the activity of receptor for adrenomedullin, calcitonin, and amylin [26–28], along with the membrane protein coding gene *KANTR*, were downregulated in both the adrenal and thyroid glands, but were upregulated in the testis of the T.poor group (**Figure 3C**). Another gene, *NPFF*, which encodes a morphine-modulating peptide, was found to be downregulated in both the adrenal and thyroid glands of the T.poor group, but showed no significant change in the testes (**Figure 3D**). These data indicate that the inflammatory state and endocrine regulation in both the adrenal and thyroid glands may be associated with the spermatogenic capacity of the testes.

### Multi-tissue transcriptomic associations with testicular function

To investigate systemic associations with spermatogenic capacity, we analyzed RNA-seq data from various somatic tissues in addition to testes, including the lung, liver, stomach, skin (sun-exposed), visceral adipose tissue, left ventricle of the heart, whole blood, tibial artery, and tibial nerve. To control for potential confounding effects of mechanical ventilation on pulmonary transcriptomes, we excluded subjects with a history of ventilator use. This stringent filtering resulted in 11 eligible subjects (6 T.good subjects and 5 T.poor subjects; **Dataset S2**). Differential expression analysis (p-value < 0.01; mean log2(TPM + 1) >0.01) revealed 6595 DE genes in the testes, 189 DE genes in the lung, 90 DE genes in the liver and fewer than 90 DE genes in each of the other tissues (**Figure 4A** and **Dataset S5**). These data suggest that the lung and liver may be more closely related to testicular function compared to the other tissues or organs.

**Figure 4.**
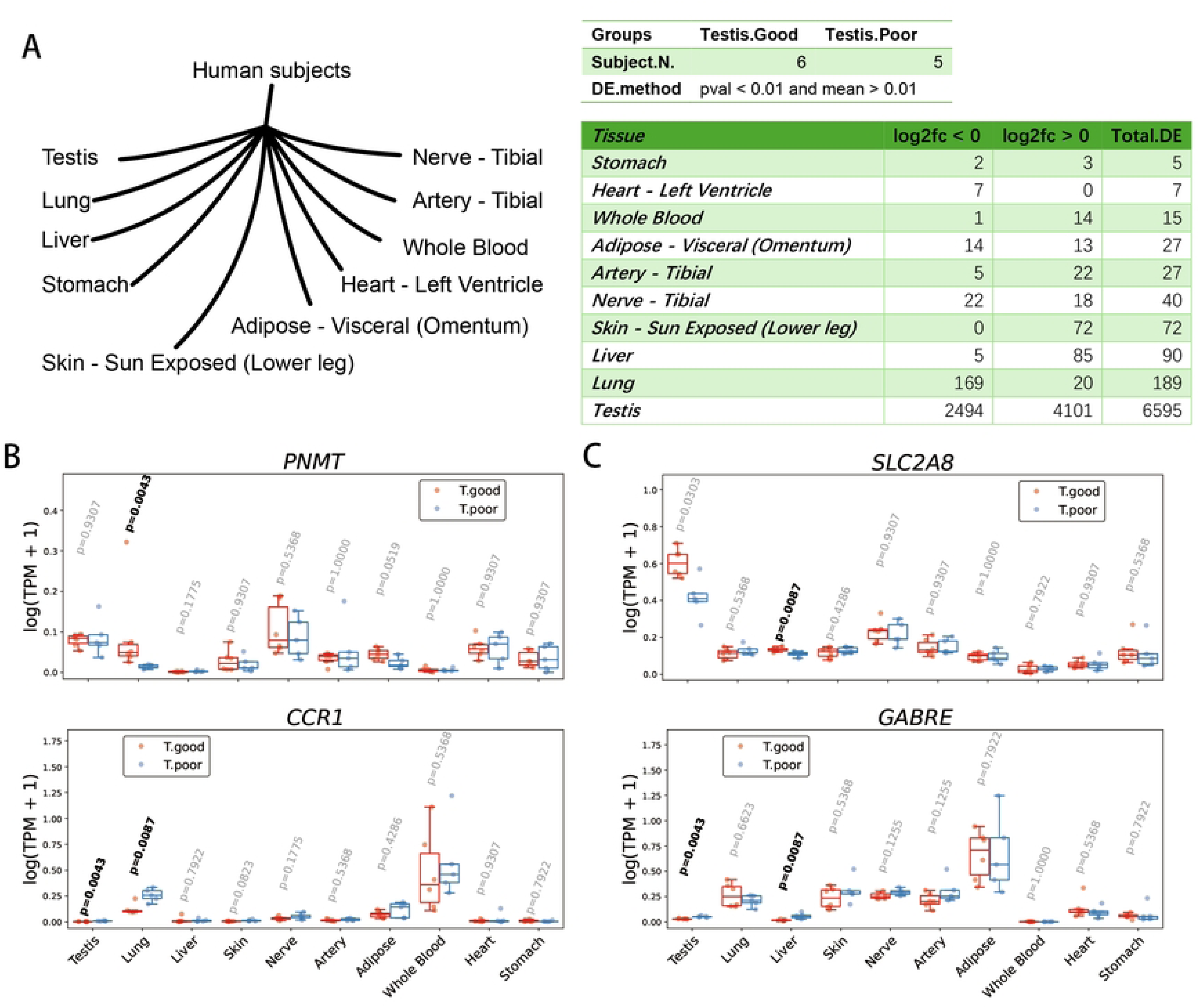
Genes in the lung and other tissues/organs associated with testicular function. (**A**) Differentially expressed (DE) genes in various tissues/organs based on differing testicular spermatogenesis abilities. (**B**) Genes that are differentially expressed in the lung. (**C**) Genes that are differentially expressed in the liver.

In the lungs of the T.poor group, the *PNMT* gene, which encodes a methyltransferase that converts nonepinephrine into epinephrine and also acts on phenylethanolamine and octopamine, was downregulated in the lungs but not in other tissues. Conversely, the *CCR1* gene, which regulates immune responses, was upregulated in both the lungs and testis (**Figure 4B**). In the liver of the T.poor group, the *SLC2A8* gene, which encodes a membrane protein responsible for the transport of glucose and fructose, was downregulated in both the liver and testis. Additionally, the *GABRE* gene, which encodes the gamma-aminobutyric acid receptor subunit epsilon, was upregulated in the liver and testis (**Figure 4C**). These data suggest that immune responses, metabolism, and endocrine regulation may be associated with the spermatogenic capacity of the testis.

### Transcriptomic associations between testicular function and multiple brain regions

Beyond the hypothalamic-pituitary axis, our study reveals significant correlations between testicular function and various brain regions involved in emotional and cognitive processing. We analyzed RNA-seq data from testes and 11 distinct brain regions in 14 human subjects (6 T.good, 8 T.poor; **Dataset S2**), identifying tissue-specific patterns of differential gene expression (p-value < 0.01; mean log2(TPM + 1) > 0.01): 5965 DE genes in the testes, 163 DE genes in the amygdala, 139 DE genes in the substantia nigra, 133 DE genes in the anterior cingulate cortex, 124 DE genes in the frontal cortex, and fewer than 100 DE genes in each of the other brain regions (**Dataset S6** and **Figure 5A**).

**Figure 5.**
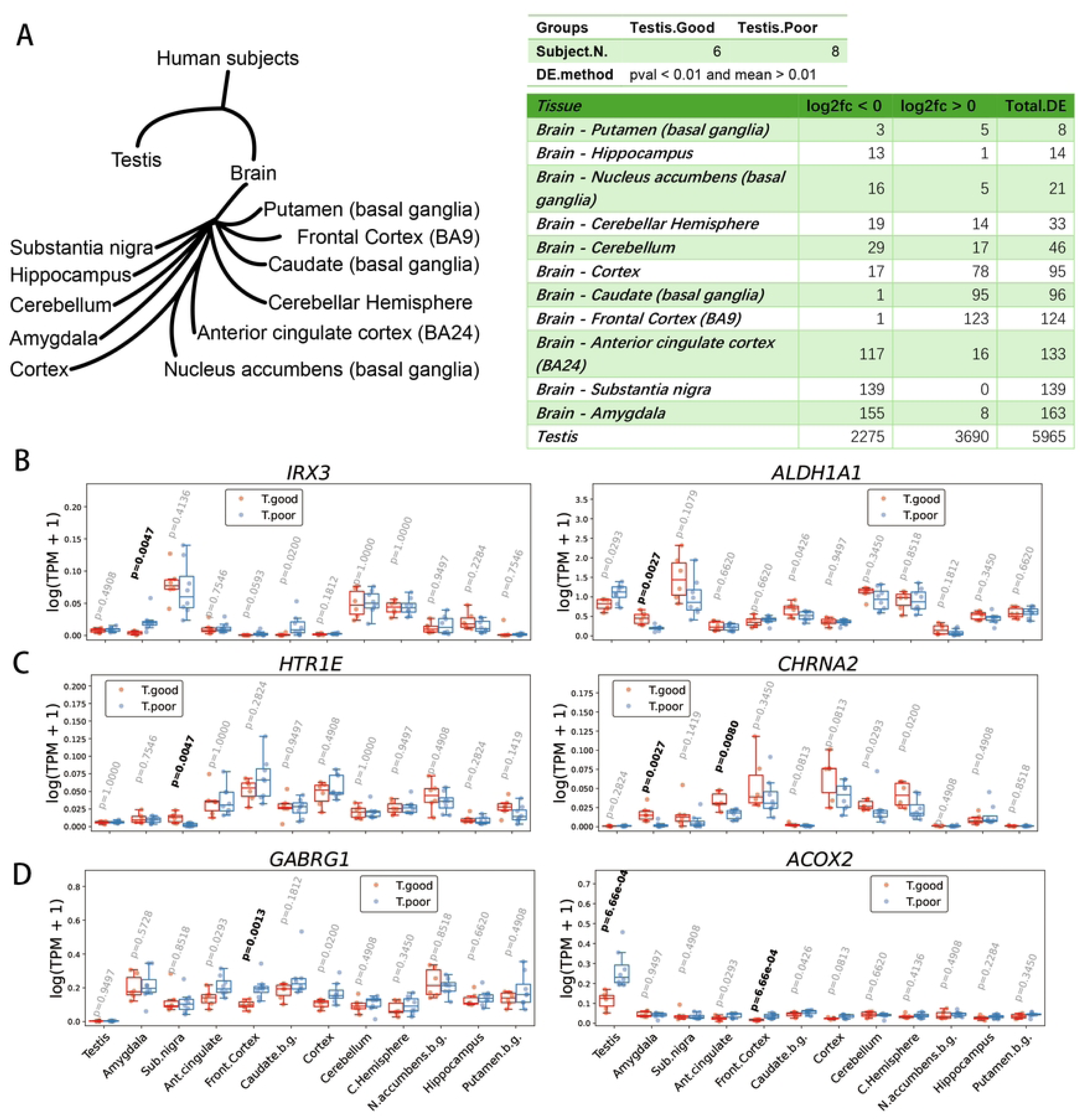
Brain regions associated with testicular spermatogenesis. (**A**) Differentially expressed (DE) genes in brain regions based on varying abilities of testicular spermatogenesis. (**B**) Genes that are differentially expressed in the amygdala. (**C**) Genes that are differentially expressed in substantia nigra. (**D**) Genes that are differentially expressed in the anterior cingulate cortex and frontal cortex.

In these brain regions, the amygdala is involved in emotional processes such as fear [29]. The substantia nigra is associated with voluntary movement and reward processing in humans [30], and the loss of dopaminergic neurons in substantia nigra may lead to Parkinson’s disease [31]. The anterior cingulate cortex and frontal cortex also play a role in regulating emotional functions [32]. In the amygdala, we found that the *IRX3* gene was upregulated in the T.poor group, while the *ALDH1A1* gene was downregulated (**Figure 5B**). The 5-hydroxytryptamine receptor gene *HTR1E* was downregulated in the substantia nigra of the T.poor group, and the neuronal acetylcholine receptor gene *CHRNA2* was downregulated in the anterior cingulate cortex and amygdala of the T.poor group (**Figure 5C**). In both the anterior cingulate cortex and frontal cortex, we found that the gamma-aminobutyric acid receptor gene *GABRG1* and the peroxisomal acyl-coenzyme A oxidase gene *ACOX2* were upregulated in the T.poor group. These data suggest that testicular function may be associated with brain regions involved in emotional processing.

In the brain, the activation, inhibition, and communication between neurons rely on neurotransmitters that function on synapses. We analyzed additional neurotransmitter receptors that may be associated with testicular function (**Dataset S7**). As a result, we found that the dopamine receptor gene *DRD4* was downregulated in the hippocampus of the T.poor group (**Figure 6A**), while the glutamate receptor gene *GRIN2D* was downregulated in the amygdala of the T.poor group (**Figure 6B**). In the substantia nigra, in addition to *HTR1E*, the acetylcholine receptor gene *CHRM4*, the glycine receptor gene *GLRB*, the gamma-aminobutyric acid receptor genes *GABRA2* and *GABRB3*, as well as the glutamate receptor genes *GRIK2* and *GRM1* were all downregulated in samples from the T.poor group (**Figure 6C**). These data indicate a significant correlation between testicular function and neuronal communication in specific brain regions, particularly in the substantia nigra.

**Figure 6.**
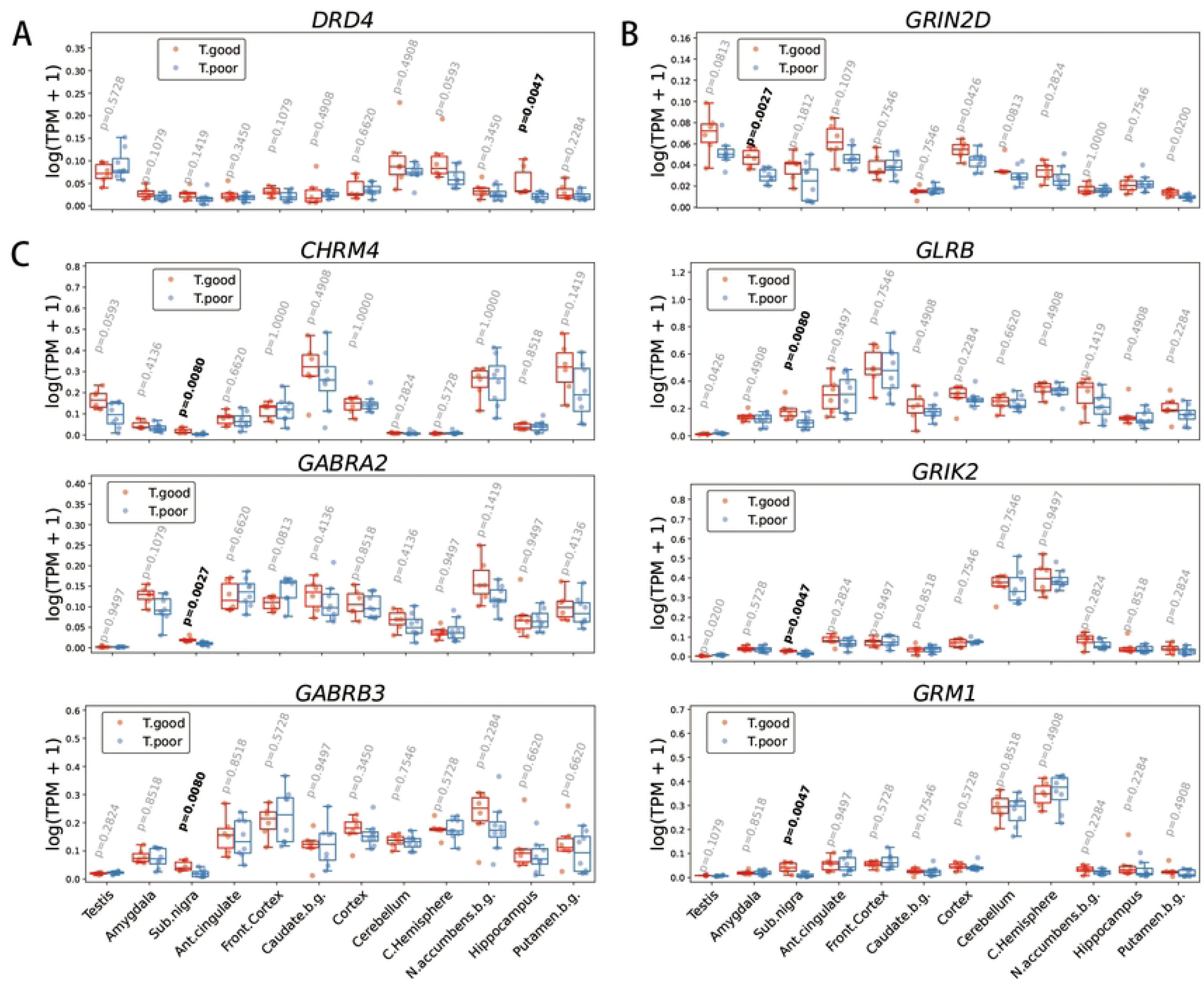
Differentially expressed brain neurotransmitter receptors associated with testis function. (**A**) The dopamine receptor gene *DRD4* was downregulated in the hippocampus of subjects in the T.poor group. (**B**) The glutamate receptor gene *GRIN2D* was downregulated in the amygdala of T.poor subjects. (**C**) Several neurotransmitter receptor genes were downregulated in the substantia nigra of T.poor subjects.

## Discussion

Understanding the relationship between testicular spermatogenic capacity and systemic tissues and organs is crucial for elucidating the interplay between reproductive health and overall physiological homeostasis. Using data from GTEx, we stratified human subjects into groups with higher and lower testicular spermatogenic capacity and analyzed the transcriptomic differences in systemic tissues and organs between the two groups. In addition to the HPT axis and adrenal gland, we found that the lung and liver exhibit a stronger association with testicular spermatogenic capacity compared to tissues such as blood and stomach. Furthermore, in the brain, regions associated with emotional regulation, including the amygdala, substantia nigra, anterior cingulate cortex, and frontal cortex, also demonstrate certain correlations with testicular function. These findings suggest that the development of drugs targeting testicular function should not only be approached from a multi-omics perspective [33–36], but also consider systemic interactions across different tissues and organs.

Male infertility can result from either monogenic variants [37] or the combined effects of environmental factors and polygenic susceptibility [2,38]. In the adrenal gland, high expression of inflammatory factors is associated with decreased spermatogenic capability in the testes. The lungs are directly exposed to airborne exogenous substances (e.g., pathogens, pollutants), making them highly susceptible to triggering immune responses [39]. The liver’s metabolic capacity is intrinsically linked to immune regulation through the detoxification of immunomodulatory substances, synthesis of complement and clotting factors, and antigen presentation by Kupffer cells and hepatocytes [40]. In testes with lower spermatogenetic capability, inflammation-associated genes such as *C3*, *C4B*, and *ATRN* are also upregulated. These findings suggest that abnormalities in the lungs and liver could elevate systemic inflammation, consequently impairing testicular function. In humans, a chronic inflammatory microenvironment can lead to various detrimental consequences, including carcinogenesis [41,42]. The pro-inflammatory polarization of tissue-resident macrophages may not only exert local effects but also remotely impact testicular function, while anti-inflammatory agents such as curcumin [43] could potentially enhance male fertility.

Lifestyle modification is considered a potential approach to addressing male infertility caused by testicular failure [44]. In this study we found that inflammation and neuroendocrine factors corelate with testicular function. The UK Biobank data has showed that environmental particulate matter pollution elevates lung inflammation [45]. Air pollution has been considered as an inconclusive risk factor of male infertility [46], therefore, we speculate that air pollution may lead to a decline in testicular function by inducing inflammatory responses in the lungs, and reducing exposure to air pollution could potentially help improve male fertility.

In the brain, the substantia nigra primarily regulates motor control and reward processing through the production of dopamine [47]. In this study, we identified a significant relationship between the testes and the differentially expressed neurotransmitter receptors in the substantia nigra. Additionally, it has been reported that testosterone can modulate the expression of dopamine-related genes in this region [48]. These findings suggest that disorders in the testes may manifest as changes in human behavior through the substantia nigra and/or other brain regions. Conversely, emotional states may significantly impact testicular spermatogenic function. Cultivating positive psychological well-being could potentially enhance sperm production, suggesting a novel adjunctive approach for male patients undergoing assisted reproductive treatments. However, further research and analysis are needed to determine whether and to what extent variations in testicular function may influence human behavioral traits.

In this study, we identified differential expression patterns of bioactive peptide-encoding genes: *NPFF* was downregulated in the adrenal gland and thyroid, while *NMS* showed downregulation and *EDN1* exhibited upregulation in the hypothalamus. These neuropeptides may serve as potential biomarkers for both detecting and investigating the underlying causes of male subfertility, or as potential therapeutic targets for indirectly improving testicular spermatogenic function.

This study has some primary limitations. First, it reveals that the relationships between the testes and various tissues or organs are correlational rather than causal. Consequently, we cannot ascertain whether changes in these tissues or organs affect testicular spermatogenic capacity, whether differences in spermatogenic capacity influence other tissues or organs, or whether an underlying abnormality simultaneously impacts both the testes and a specific tissue or organ. Second, the sample size per group is limited when comparing the strength of associations between various tissues or organs and testicular function, which may increase the potential errors due to sampling bias. Third, the GTEx version 10 dataset, collected from postmortem donor specimens, may yield under- or overestimated functional assessments of specific genes in certain tissues or organs due to potential inconsistencies in RNA-seq protocols, including tissue procurement, RNA extraction, and library preparation for sequencing, across different individuals. Finally, it should be noted that the observed mRNA expression differences in this study may not necessarily reflect corresponding protein levels in tissues, and since individual genetic variations were not incorporated into our analysis, the upregulation or downregulation of mRNA alone may not accurately represent functional changes in protein activity.

## Data availability

The GTEx V10 open access data (https://www.gtexportal.org/) were used in this study. The scripts used in this research have been uploaded to GitHub (https://github.com/MaJYatGZ/GTEx_testis_Analysis).

## Acknowledgements

The Genotype-Tissue Expression (GTEx) Project was supported by the Common Fund of the Office of the Director of the National Institutes of Health, and by NCI, NHGRI, NHLBI, NIDA, NIMH, and NINDS. The data used for the analyses described in this manuscript were obtained from the GTEx Portal (https://www.gtexportal.org/) on 03/10/2025. We thank the staff at the GD2H Reproductive Medicine Center for their supports and assistances in this study.

## Authors’ roles

JYM, JC and YC contributed the central idea; JYM and XA analyzed most of the data; JYM, XA, FYX, JC, and YC wrote the paper; All authors contributed to refining the ideas and finalizing this paper.

